# Integrating deep mutational scanning and low-throughput mutagenesis data to predict the impact of amino acid variants

**DOI:** 10.1101/2022.12.14.520494

**Authors:** Yunfan Fu, Justin Bedő, Anthony T. Papenfuss, Alan F. Rubin

## Abstract

Evaluating the impact of amino acid variants has been a critical challenge for studying protein function and interpreting genomic data. High-throughput experimental methods like deep mutational scanning (DMS) can measure the effect of large numbers of variants in a target protein, but because DMS studies have not been performed on all proteins, researchers also model DMS data computationally to estimate variant impacts by predictors. In this study, we extended a linear regression-based predictor to explore whether incorporating data from alanine scanning (AS), a widely-used low-throughput mutagenesis method, would improve prediction results. To evaluate our model, we collected 146 AS datasets, mapping to 54 DMS datasets across 22 distinct proteins. We show that improved model performance depends on the compatibility of the DMS and AS assays, and the scale of improvement is closely related to the correlation between DMS and AS results.

## 1 Introduction

Deep mutational scanning (DMS) is a functional genomics method that can experimentally measure the impact of many thousands of protein variants by combining high-throughput sequencing with a functional assay (Fowler & Fields, 2014). In a typical DMS, a cDNA library of genetic variants of a target gene is generated, containing all possible single amino acid substitutions. This variant library is then expressed in a functional assay system where the variants can be selected based on their properties. The change in variant frequency in the pre- and post-selection populations is determined by high-throughput sequencing which is then used to calculate a multiplexed functional score that captures the variant’s impact (Findlay, 2021; Geck et al., 2022; Weile & Roth, 2018). The versatility of DMS assays makes it possible to measure variant impact on a wide range of protein properties, including protein binding (Diss & Lehner, 2018; Fowler et al., 2010), protein abundance (Amorosi et al., 2021; Faure et al., 2022; Matreyek et al., 2018), catalytic activity (Mighell et al., 2018; Stiffler et al., 2015) and cell growth rate (Ahler et al., 2019; Giacomelli et al., 2018; Roscoe et al., 2013).

Computational studies have used DMS data to build predictive models of variant impact. These predictors use supervised or semi-supervised learning models trained on experimental DMS data and various protein features to make predictions (Gray et al., 2018; Alley et al., 2019; Munro & Singh, 2020; Biswas et al., 2021; Høie et al., 2022; Wu et al., 2021; Hsu et al., 2022). Envision is one such method that used protein structural, physicochemical, and evolutionary features to predict variant effect scores and was trained on DMS data from 8 proteins using gradient boosting (Gray et al., 2018). Another method, DeMaSk, predicted DMS scores by combining two evolutionary features (protein positional conservation and variant homologous frequency) with a DMS substitution matrix and was trained on data from 17 proteins using a linear model (Munro & Singh, 2020). Deep learning algorithms have also been applied to build protein fitness predictors (Alley et al., 2019; Biswas et al., 2021), which are usually based only on variant sequences.

Low-throughput mutagenesis experiments that measure tens of variants at a time have also been used extensively to study diverse protein properties, including substrate binding affinity (Block et al., 1996; Sloan & Hellinga, 1999), protein stability (Fleming & Engelman, 2001; Shibata et al., 2009), and protein activity (Brzovic et al., 2011; Gajula et al., 2014). Alanine scanning (AS) is a widely-used low-throughput mutagenesis method (Kortemme et al., 2004; Morrison & Weiss, 2001), and AS data are available for many proteins. In this method, each targeted protein residue is substituted with alanine, and the impacts of these variants are measured by a functional assay (Cunningham & Wells, 1989). AS experiments are typically used to identify functional hot spots or critical residues in the target protein (DeLano, 2002; Eustache et al., 2016) and have been used as a source of independent validation for DMS studies (Gajula et al., 2014; Olson et al., 2014; Staller et al., 2018; Gray et al., 2019).

In this study, we explore whether a predictive model can be improved by incorporating low-throughput mutagenesis data (Fig 1). We find that AS data can increase prediction accuracy and that the improvement is related to the similarity of the functional assays and the correlation of DMS and AS results.

**Fig 1.**
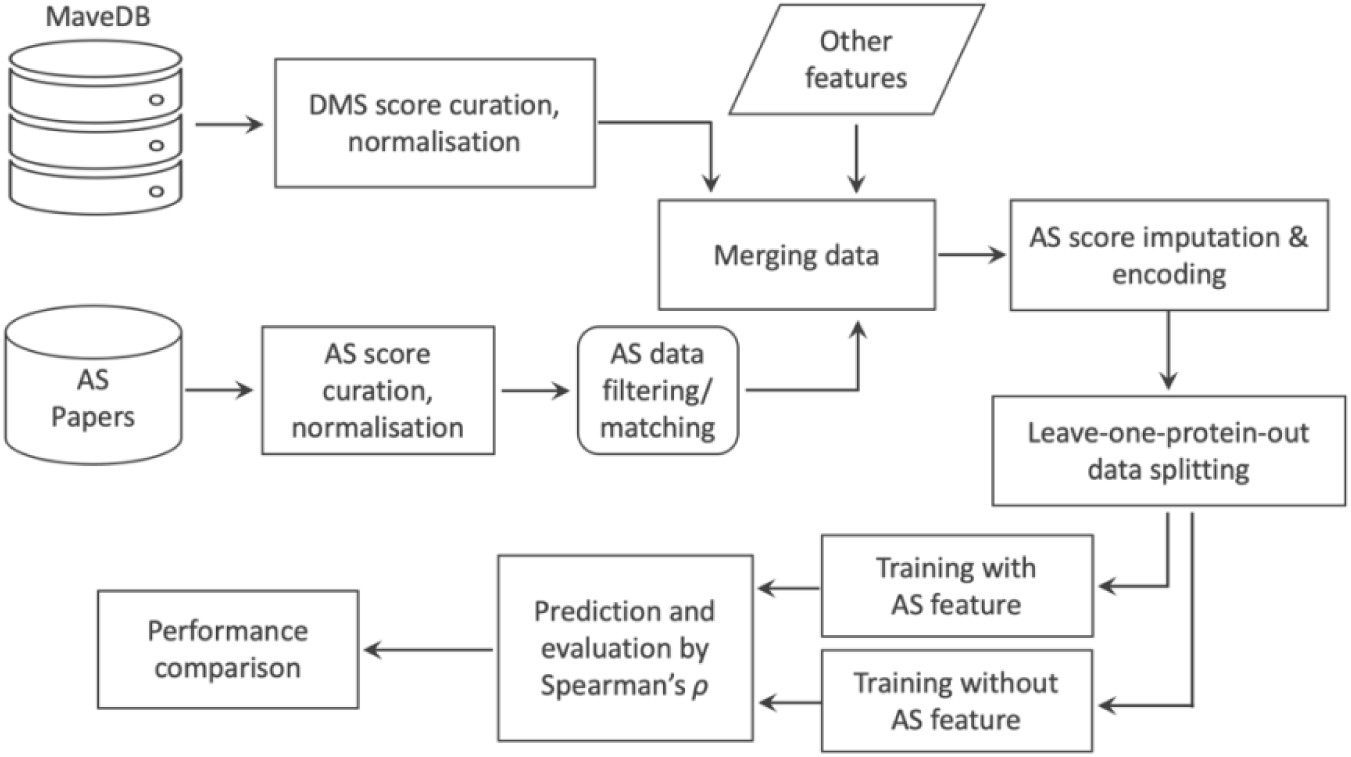
Workflow for model training and testing. DMS and AS datasets are collected from online resources and are normalized. DMS and AS datasets targeting the same protein are then matched, filtered and merged. Two predictors are constructed and tested: the first uses DMS data, AS data and other protein features, and the second uses only DMS data and the same other protein features.

## 2 Results

### 2.1 Overview of DMS and alanine scanning (AS) data

To build the predictive model, 130 DMS datasets were collected from MaveDB (Esposito et al., 2019; Rubin et al., 2021) (Supplementary table 1). We searched the literature and found 146 AS datasets targeting the same proteins as 54 of the DMS datasets. In total, we obtained both DMS and AS data for 22 different proteins: 17 human proteins, three yeast proteins, and two bacterial proteins. Most DMS experiments were highly complete, with a mean coverage of 95.0% of all possible single amino acid substitutions assayed in the target region, comprising 373,219 total protein variant measurements. AS data were only available on a small number of protein residues (Fig 2), and we were able to curate 1,480 alanine substitution scores from the 146 studies. Variant scores from collected DMS and AS studies were linearly normalized to a common scale (Gray et al., 2018, see Methods) to make them comparable across datasets (Fig S1).

**Fig 2.**
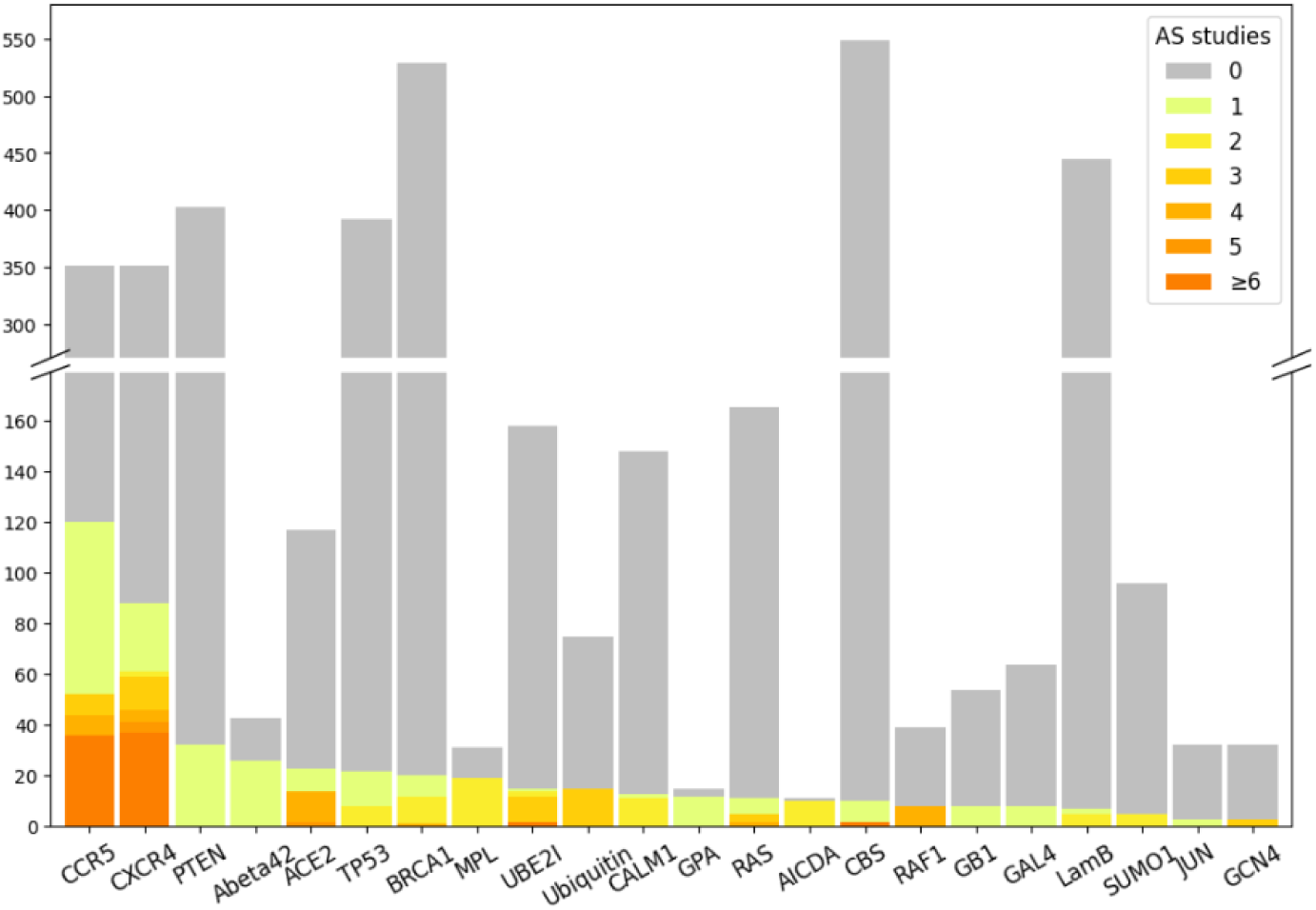
DMS data generally cover more protein residues than AS data. Each bar shows the number of residues assayed by DMS studies on given target proteins. Colour indicates the number of AS studies available for the DMS-tested residues.

### 2.2 The correlation of DMS and AS scores is related to assay compatibility

To evaluate the similarity of AS and DMS scores, we calculated Spearman’s correlation (*ρ*) between the AS scores and DMS scores for the same alanine substitutions. Since each protein may have results from several AS and DMS experiments, we calculated *ρ* between each possible pair. The median *ρ* over DMS and AS data (DMS/AS) pairs was 0.2, indicating that the experimental scores were poorly correlated overall (Fig 3).

**Fig 3.**
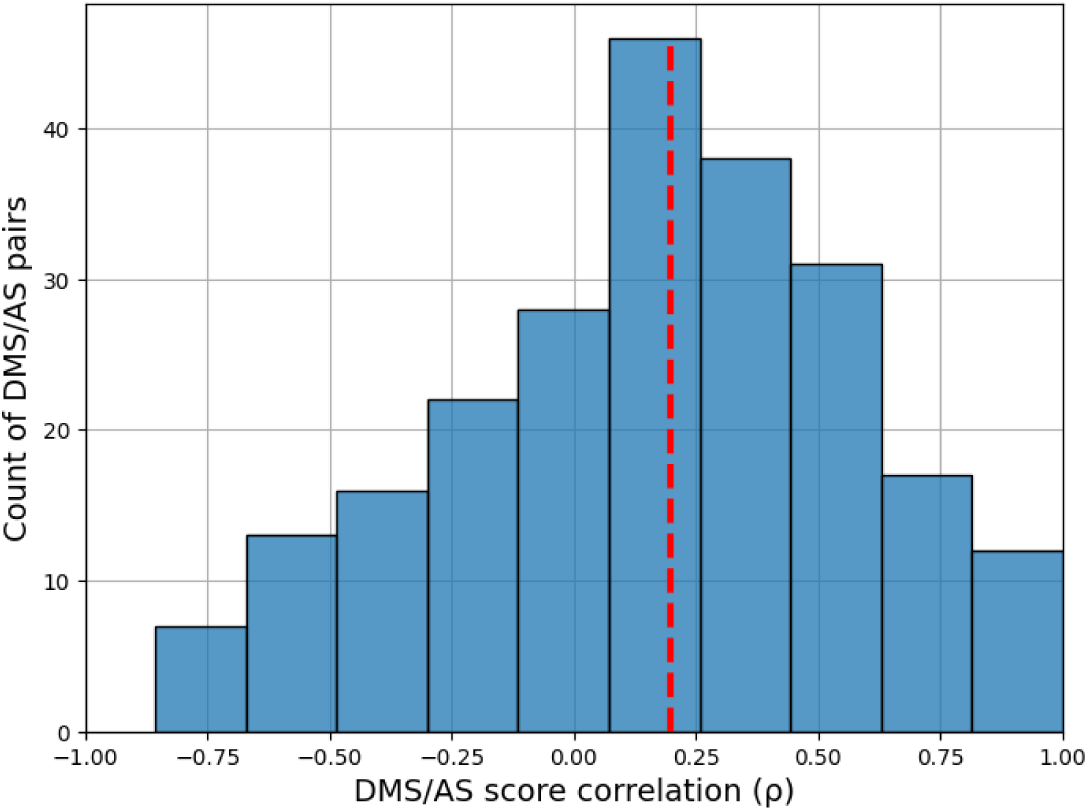
Correlation between DMS and AS data shows substantial variation. We calculated Spearman’s *ρ* between alanine substitution scores in each pair of AS and DMS data. The results for pairs with less than three alanine substitutions are removed. The red dashed line shows the median *ρ*.

We then considered if differences between AS and DMS assay designs might contribute to this low agreement between scores. To explore this, we developed a decision tree (Fig S2) to classify whether DMS/AS pairs had low, medium, or high assay compatibility, which we defined as a similarity measurement of the functional assays performed. For example, the DMS assay measuring the binding affinity of a cell surface protein, CXCR4, to its natural ligand (Heredia et al., 2018) has high compatibility with the AS experiment also measuring this ligand binding but has low compatibility with the study on CXCR4’s ability to facilitate virus infection (Tian et al., 2005). A full assay compatibility table can be found in Supplementary Table 1 with the compatibility classifications and justification for each pair. We then compared DMS and AS score correlation for each compatibility class and found that score correlations were closely related to assay compatibility. Data from low compatibility assays had a median correlation of 0.15, rising to 0.19 for medium compatibility assays and 0.40 for high compatibility assays (Fig 4). This link between assay compatibility and score correlation indicates that our decision tree approach was able to capture the similarity between assay systems.

**Fig 4.**
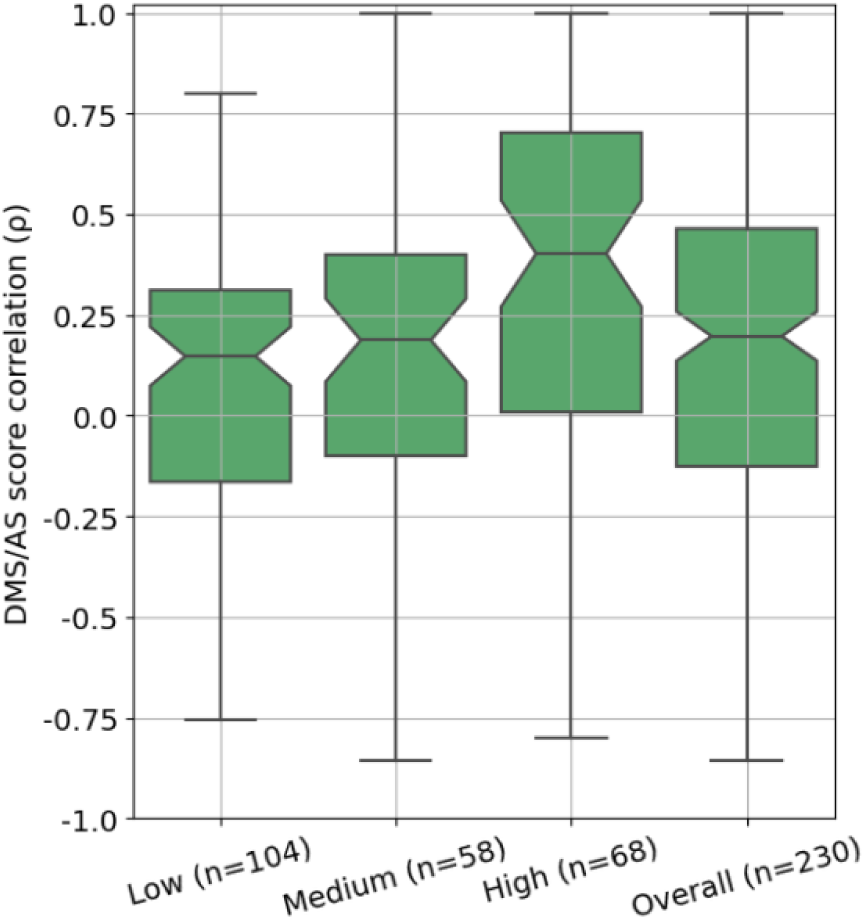
DMS and AS data pairs with high assay compatibility show a higher score correlation. Each box represents Spearman’s *ρ* between DMS and AS data pairs of classified assay compatibility or the overall result. The correlation coefficients are calculated between alanine substitution scores in each pair of AS and DMS data. Results for data pairs with less than three alanine substitutions are removed.

### 2.3 Compatible AS data improve DMS score prediction accuracy

To test if incorporating AS data into DMS score models would improve prediction accuracy, we decided to build a new model based on DeMaSk. We chose DeMaSk because it showed better performance compared to similar methods and was straightforward to modify. The published DeMaSk model predicts DMS scores using protein positional conservation, variant homologous frequency, and substitution score matrix, and we incorporated AS data as an additional feature. Our new predictor was modelled with all 130 DMS we collected and we applied a leave-one-protein-out cross-validation approach to training and testing (Gray et al., 2018). Prediction performance was evaluated using the Spearman’s correlation (*ρ*) between the experimentally-derived DMS scores and the predicted scores for each pair of DMS and AS studies. The performance of our DMS/AS model was compared with a model trained only on DMS data, equivalent to retrained DeMaSk (Fig S3), by calculating the change of prediction *ρ* (see Methods).

We trained our model with either all or a subset of AS data we collected (Fig 5, Table S1). We first integrated all 146 AS data collected for training and evaluation but observed only a modest improvement of prediction *ρ* (Fig 5 left box, and Fig S4). We then retrained and evaluated our model on filtered AS data with only high compatibility assays, and observed a median increase in prediction Spearman’s *ρ* of 0.1 compared to the results with no AS data (Fig 5 middle box, and Fig S4). However, training with both high and medium compatibility pairs reduced the performance improvement (Fig S5). These results indicate that low and medium compatibility pairs might provide inconsistent training data, degrading model performance. We also evaluated the impact of including high compatibility AS data in an alternative model based on Envison (Gray et al., 2018), and found similar results (Fig S6). To differentiate between high assay compatibility and high DMS/AS score correlation, we trained the model using the most highly correlated AS result for each DMS dataset (see Methods). Although the upper quartile was high, the median performance change of this predictor was lower than the high assay compatibility model, suggesting that matching with the highest score correlation alone is insufficient (Fig 5 right box).

**Fig 5.**
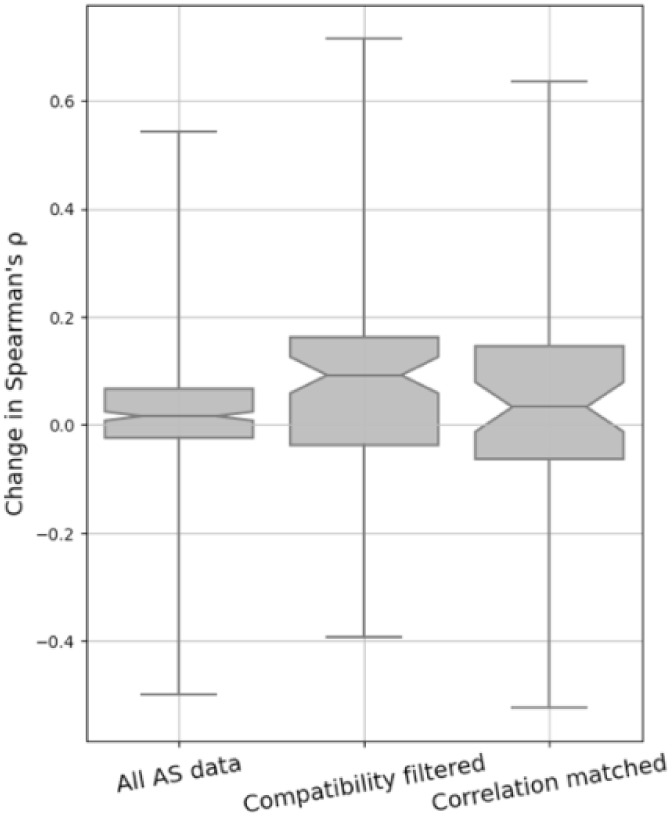
Performance of variant impact prediction is improved using AS data with high assay compatibility. The change of prediction *ρ* for each DMS and AS data pair is shown as box plots. A higher value represents higher prediction accuracy achieved for using AS data. Different approaches to filtering/matching the data are shown on the x-axis: “All AS data” used all available data; “Compatibility filtered” used only data of high assay compatibility; “Correlation matched” used only data with the highest regularised correlation for each DMS dataset.

To further explore the higher performance of compatibility-filtered predictor, we examined the relationship between prediction *ρ* change and score correlation for each high compatibility DMS/AS pair (Fig 6). For most pairs, prediction performance was improved by using AS data, and the scale of improvement was also related to the score correlation. This relationship could also be observed for multiple DMS/AS pairs from an individual protein, such as CXCR4 and CCR5. We saw the same trend in the predictor trained with all DMS/AS pairs but noted that the performance even of highly correlated pairs was worse, likely due to the influence of low compatibility training data on the model (Fig S7).

**Fig 6.**
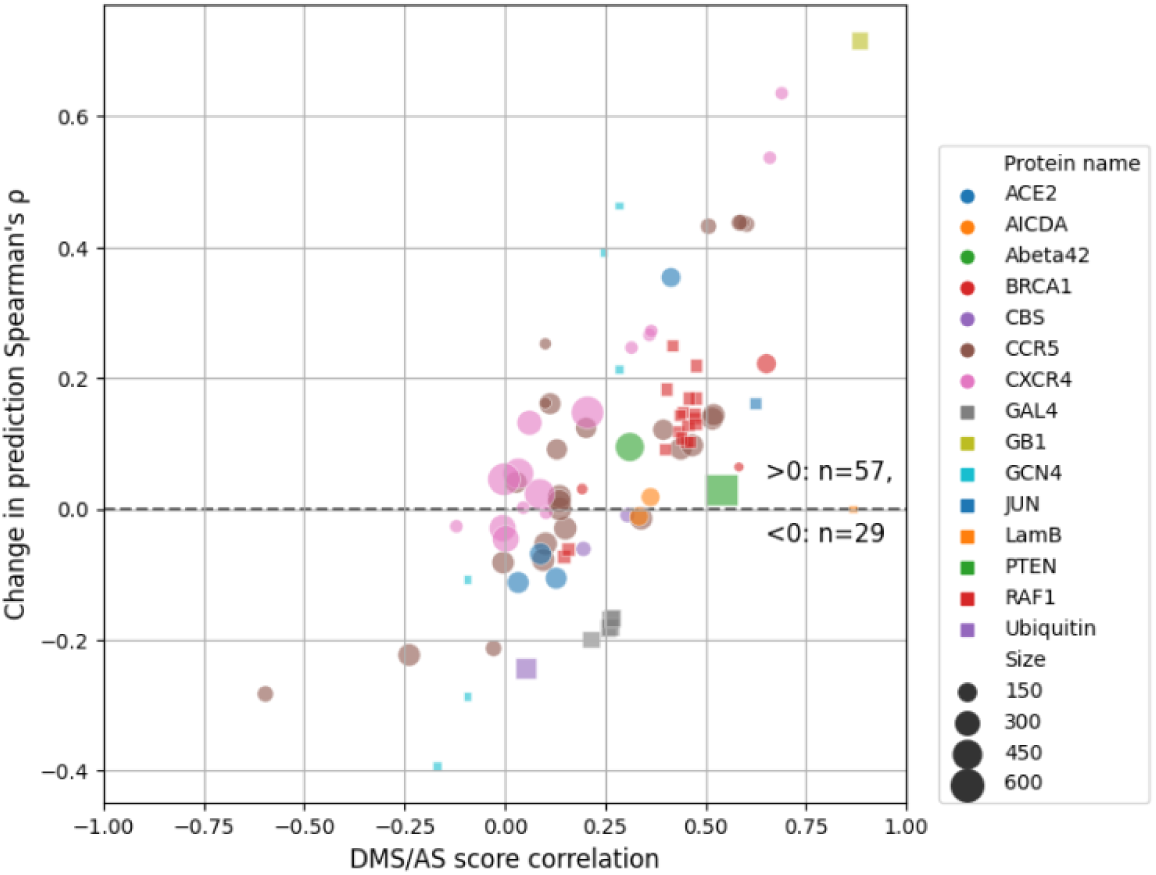
Prediction performance change is related to DMS and AS score correlation. Each dot represents a filtered DMS/AS data pair of high assay compatibility. The vertical axis shows the change of prediction *ρ* by using AS data (larger means higher performance achieved by using AS data). The horizontal axis shows the DMS/AS score correlation for *all* variants on the matched residues rather than just alanine substitutions. The colours and shapes of the dots correspond to the target protein, and size indicates the number of variants in each data pair.

We also explored the consequences of the sparsity of AS data on our model in two ways: by using a boosting approach that focuses only on residues with AS data (Fig S8) and by using complete alanine substitution information from DMS as the AS feature (Fig S9). Both of these approaches performed very similarly to the primary model constructed using high-compatibility DMS/AS data and simple mean score imputation.

To test the influence of amino acids on our predictor, we grouped the prediction results by either wild-type or variant amino acid and calculated the prediction improvement when AS data were included (Fig 7). We found that 14 of 19 wild-type amino acids performed better with the addition of AS data, with cysteine showing the largest improvement and performing worst in the model lacking AS data. 18 of 20 variant amino acids benefited from the inclusion of AS data, with marginal performance decrease on lysine and aspartic acid (|Δ*ρ*|<0.01) (Fig 7).

**Fig 7.**
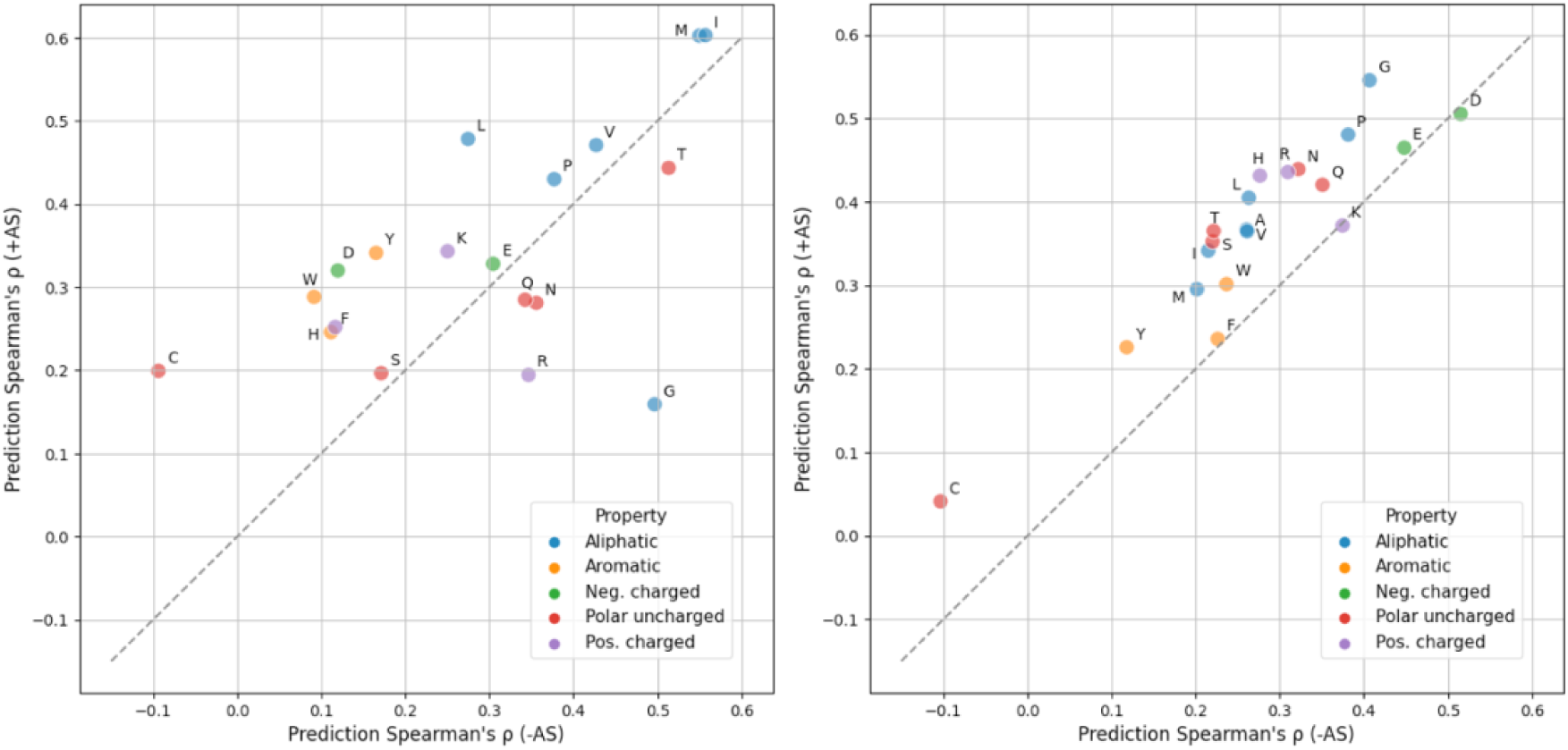
Model perfomance is generally improved for each wild-type and variant amino acid. Prediction Spearman’s *ρ* when using (y-axis) or not using (x-axis) AS data on each wild-type (left) or variant (right) amino acid is shown in the scatter plots. The results are coloured according to the property of each amino acid type. Alanine (A) result is not applicable in the first figure since alanine scanning data are always missing when the wildtype is alanine itself. Absolute count for each amino acid can be found in Fig S10. (Neg.: negatively, Pos.: positively)

## 3 Discussion

In this study, we integrated alanine scanning (AS) data into deep mutational scanning (DMS) score prediction, leading to modest improvements in the accuracy of variant score prediction. We also explored the impact of the diversity of protein properties measured by DMS and AS. Filtering DMS and AS data based on our manual classification of assay type compatibility led to improved prediction performance.

A potential shortcoming of our current approach is that AS data were available for only a small proportion of the DMS data. Although most recent DMS studies can analyze variants of the whole protein, most AS experiments only cover a handful of residues in the target protein, leaving missing AS scores for the vast majority of residues. We explored this here and found that alternative methods for addressing the sparsity of AS data did not improve or degrade performance, but we anticipate further improved prediction accuracy if the low completeness and unevenness of AS data are appropriately handled before modelling, such as by advanced imputation methods (Stekhoven & Buhlmann, 2012; Wu et al., 2019).

In this study, we identified the importance of DMS/AS assay compatibility as a crucial factor for improving prediction accuracy. An issue with using this concept is that it further shrinks already sparse data. It also fails to take advantage of the fact that even for low compatible assays some fundamental information like protein stability can still be mutually captured. Instead of hard filtering, proper implementation of this underlying information may facilitate variant impact prediction in the future. Nonetheless, filtering on assay compatibility still leads to performance improvement. We also briefly explored whether the consistency of DMS and AS scores can be considered more directly by matching the best correlated AS data for each DMS dataset. Consistency is partially driven by assay compatibility but also reflects other features of the data, such as bias and noise. While we picked the most correlated pair for each DMS, we did not threshold the correlation, potentially including data pairs that were poor matches.

The concepts of compatibility and data quality are also relevant to training any DMS-based predictors. DMS assays have been developed to measure variant impacts to distinct protein properties, and a variant can behave similarly to wildtype when measured by one assay yet show altered protein properties in other assay results, which are frequently found in regions with specific biochemical functions (Cagiada et al., 2021, 2022; Jepsen et al., 2020; Matreyek et al., 2021; Mighell et al., 2020; Nielsen et al., 2021). With more experimental assays to be applied, the diverse measurements may impede the progress of future DMS-based predictors unless this assay effect is properly addressed, for example, by building assay specific predictors. Measurement error is another source of DMS data heterogeneity that potentially affects the model performance. In our current study, DMS scores of protein variants are weighted equally while training. Adjustable weighting can be applied in future studies to adapt the distinct experimental error between individual variants and datasets, reducing the influence of low-confident data.

In summary, we conclude that the careful inclusion of low-throughput mutagenesis data improves the prediction of DMS scores, and the approaches described here can potentially be applied to other prediction methods.

## 4 Supporting information

The code of this study is available at: https://github.com/PapenfussLab/DMS_with_Alanine_scan

**Supplementary Table 1:** All candidate DMS and alanine data with detailed dataset information.

**Supplementary Table 2:** Normalized DMS dataset with protein property features.

**Supplementary Table 3:** Normalized alanine scanning dataset.

## 5 Author contributions

YF developed the software and wrote the initial draft of the manuscript. AFR conceived the study. JB, AFR, and ATP oversaw the project. All authors reviewed, contributed to, and approved the manuscript.

## 7 Methods

### 7.1 DMS data collection

DMS data were downloaded from MaveDB (Esposito et al., 2019; Rubin et al., 2021) which were then filtered and curated. DMS experiments targeting antibody and virus proteins were removed because of their potentially unique functionality. We retrieved the UniProt accession ID of target proteins by searching the protein names or sequences in UniProt (The UniProt Consortium et al., 2021), and proteins lacking available UniProt ID were also excluded. Datasets that are computationally processed or their wildtype-like and nonsense-like scores (see Normalization) cannot be identified were also filtered out (Supplementary Table 1). All missense variants with only a single amino acid substitution were curated from the DMS studies for our analysis. A total of 130 DMS experiments from 53 studies (Ahler et al., 2019; Andrews & Fields, 2020; Bandaru et al., 2017; Bolognesi et al., 2019; Bridgford et al., 2020; Chan et al., 2020; Chiasson et al., 2020; Diss & Lehner, 2018; Elazar et al., 2016; Findlay et al., 2018; Firnberg et al., 2014; Fowler et al., 2010; Gajula et al., 2014; Giacomelli et al., 2018; Gray et al., 2019; Heredia et al., 2018; Hietpas et al., 2011, 2013; L. Jiang et al., 2013; R. J. Jiang, 2019; Keskin et al., 2017; Kitzman et al., 2015; Kotler et al., 2018; Kowalsky & Whitehead, 2016; Matreyek et al., 2018; McLaughlin Jr et al., 2012; Melamed et al., 2013; Mighell et al., 2018; Mishra et al., 2016; Nedrud et al., 2021; Newberry, Arhar, et al., 2020; Newberry, Leong, et al., 2020; Olson et al., 2014; Roscoe et al., 2013; Roscoe & Bolon, 2014; Sarkisyan et al., 2016; Silverstein et al., 2021; Staller et al., 2018; Starita et al., 2013, 2015, 2018; Stiffler et al., 2015; Suiter et al., 2020; Sun et al., 2020; Thompson et al., 2020; Trenker et al., 2021; Weile et al., 2017, 2021; Wrenbeck et al., 2019; L. Zhang et al., 2020; Zinkus-Boltz et al., 2019, two unpublished) were collected for our analysis.

### 7.2 Collection of AS data and other features

The following process was used to search for candidate AS studies. Papers were identified by searching on PubMed and Google Scholar for the “alanine scan” or “alanine scanning” together with the name of candidate proteins. While searching in Google Scholar, we included the protein’s UniProt ID rather than molecule name as the search term to reduce false positives. Appropriate AS data were collected from the search results. Western blot results were transformed to values by ImageJ if it was the only experimental data available in the study. A total 146 AS experiments were collected from 45 distinct studies (Bernier-Villamor et al., 2002; Blanpain et al., 1999; Block et al., 1996; Brzovic et al., 2003, 2011; Chabot et al., 1999; Chen et al., 2019; Chupreta et al., 2005; Cobb & Roberts, 2000; Coyne et al., 2004; Denker et al., 2005; Dragic et al., 1998, 2000; Ecsédi et al., 2020; Fleming & Engelman, 2001; Fujita– Yoshigaki et al., 1995; Gajula et al., 2014; Han et al., 2006; Hidalgo et al., 2001; Kopecká et al., 2011; Kožich et al., 2010; Kruger et al., 2003; Lee et al., 2014; Li et al., 2005; Lin et al., 2003; Mascle et al., 2013; Matthews et al., 2011; Mayfield et al., 2012; Navenot et al., 2001; Peng et al., 2016; Peterson et al., 1996; Rabut et al., 1998; Ransburgh et al., 2010; Rodríguez-Escudero et al., 2011; Sloan & Hellinga, 1999; Starita et al., 2015; Tan et al., 2017; Tian et al., 2005; Towler et al., 2013; Trent et al., 2003; Van Gelder et al., 2002; VanBerkum & Means, 1991; Wei et al., 2010; Williams et al., 2006; J. Zhang et al., 2007).

Protein features of Shannon entropy and the logarithm of variant amino acid frequency were downloaded from the DeMaSk online toolkit (Munro & Singh, 2020). The substitution score matrix feature was calculated from the mean of training DMS scores for each of the 380 possible amino acid substitutions before each iteration of cross-validation.

### 7.3 Normalization

DMS and AS datasets were normalized to a common scale using the following approach adapted from previous studies (Gray et al., 2017, 2018). Let *D* denotes a protein study measuring scores 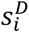 for a single variant *i*, 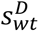 denotes the scores for wildtype and 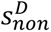 represents the score for nonsense-like variants. The normalized scores 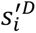 are given by:

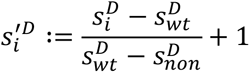

Wild-type scores were directly identified from the paper or the median score of synonymous variants. For DMS data, since not all DMS studies report score of nonsense variants, we defined the nonsense-like scores as the median DMS scores for the 1% missense variants with the strongest loss of function for each dataset. For AS data, nonsense-like scores were either defined according to the paper or using the extreme values (Supplementary Table 1).

### 7.4 AS data filtering and matching

AS data subsets were filtered/matched according to either assay compatibility or score correlation. For assay compatibility filtering, DMS and AS assay pairs were first classified into three levels of compatibility (Fig S2). For each DMS dataset, we first tried to use only AS data with high assay compatibility for further modelling, removing AS data of medium and low assay compatibility. We then also tried to model with AS data of both high and medium assay compatibility.

For score correlation matching, Spearman’s correlation (*ρ*) is calculated between alanine substitution scores in each pair of AS and DMS data. To avoid influence from the size of AS datasets, we regularised the *ρ* value by empirical copula (Nelsen, 2006):

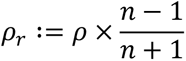

where *ρ*_r_ is the regularised correlation coefficient, and *n* is the number of alanine substitutions used for correlation calculation. For each DMS dataset, AS result with the highest *ρ_r_* was picked for modelling.

### 7.5 AS data pre-processing

AS data were pre-processed prior to modelling. For variants without available (filtered/matched) AS data, their AS scores were imputed with the mean value of all available AS scores. Then the AS data were encoded by the wild-type and variant amino acid type with one-hot-encoding. For each variant, the AS feature is expanded with two one-hot vectors. Each of the vectors has 19 zeros and one non-zero value which was the AS score, with the location of the non-zero value indicating the wild-type or variant amino acid type.

### 7.6 Training and evaluation of DMS score predictor

To build the predictors, we performed linear regression using the function sklearn.linear_model.LinearRegression from scikit-learn (Pedregosa et al., 2011). Training and validation data were separated with leave-one-protein-out cross-validation. In this process, data from one protein were withheld for subsequent validation, and the rest were used for training. This process was iterated over all proteins in the data. Variants were inversely weighted during the training process by the number of measurements available, thus compensating for some regions having greater coverage with DMS and AS assays. Predictors were trained on protein features, DMS data and (optionally) AS data using four different filtering or matching strategies: i) all DMS/AS data, ii) compatibility-filtered DMS/AS data, iii) correlation-matched DMS/AS data, and iv) a control, constructed using DMS data only. In the evaluation process, let *V* be protein variants assayed by both DMS study *D* and AS study *A*. Variant scores are predicted by the previously mentioned predictors either using AS data 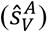 or not (*ŝ*_V_). Spearman’s correlation (*ρ*) was calculated between the DMS scores 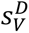 and each set of predicted scores. The difference of *ρ* was used to evaluate the performance change (Δ*ρ_v_*).

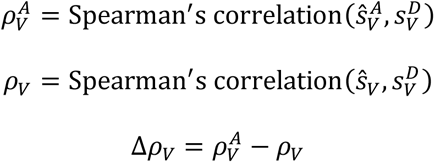

To evaluate, we iterated over variants from each pair of DMS/AS studies. Results were dropped for variants *V* with only one protein residue available during analysis and visualization.

---

**Fig S1.**
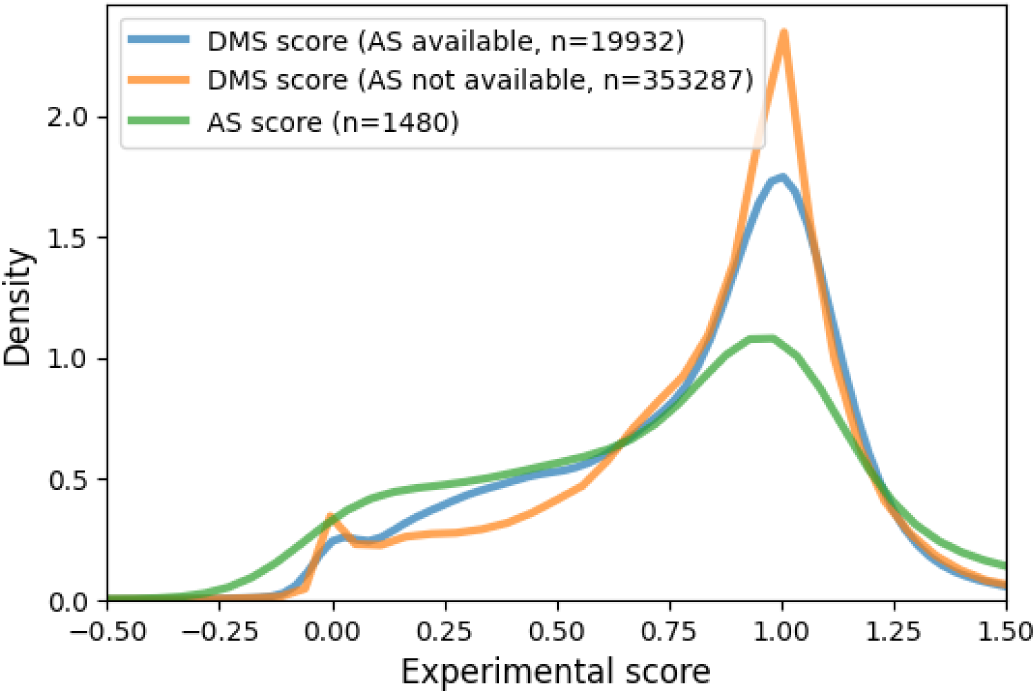
DMS and AS score distribution. The figure shows the kernel estimated density of normalized AS scores and DMS scores for variants with or without available AS data.

**Fig S2.**
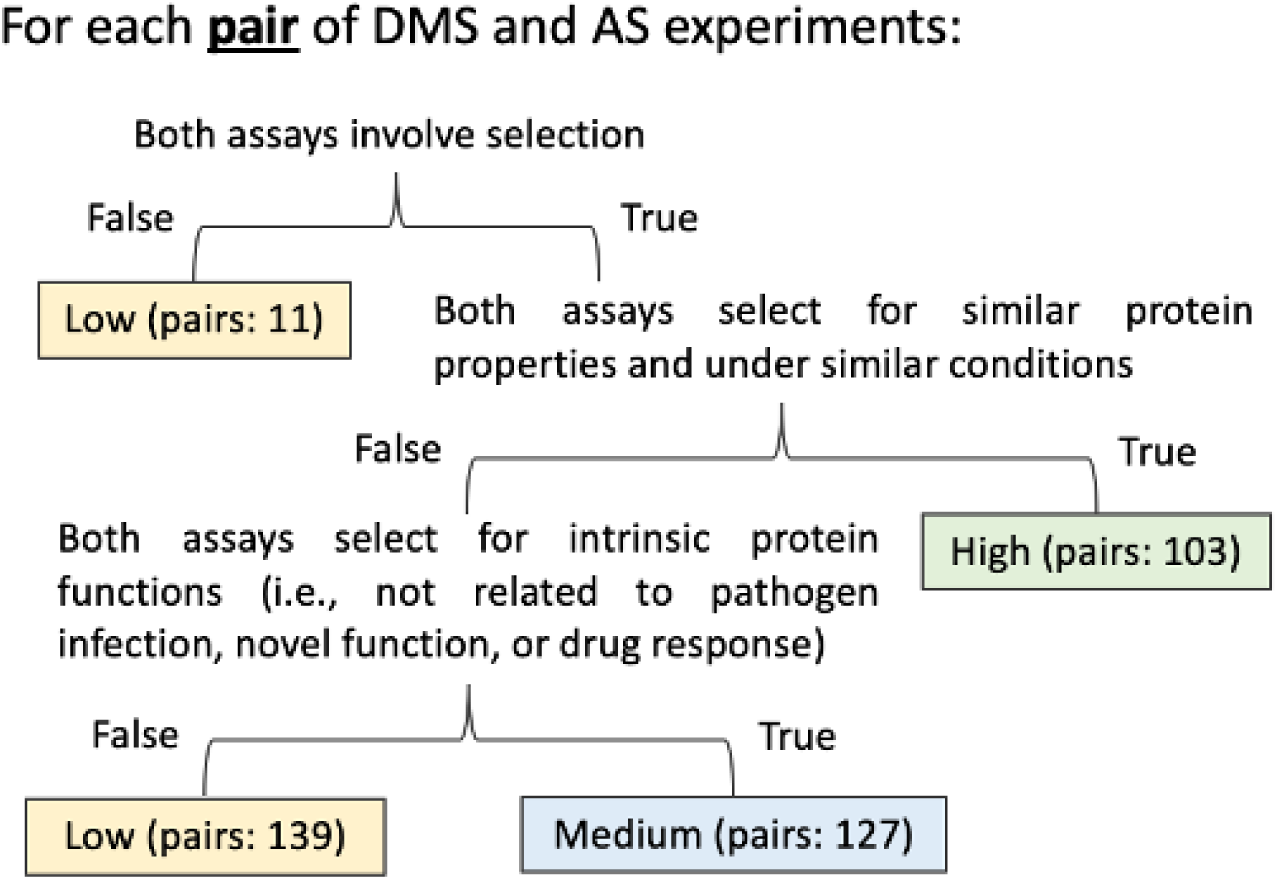
Decision tree for classifying the DMS and AS assay compatibility. The end-nodes show the classified assay compatibility. The number indicates the count of assay pairs for each compatibility level (low, medium, high).

**Fig S3.**
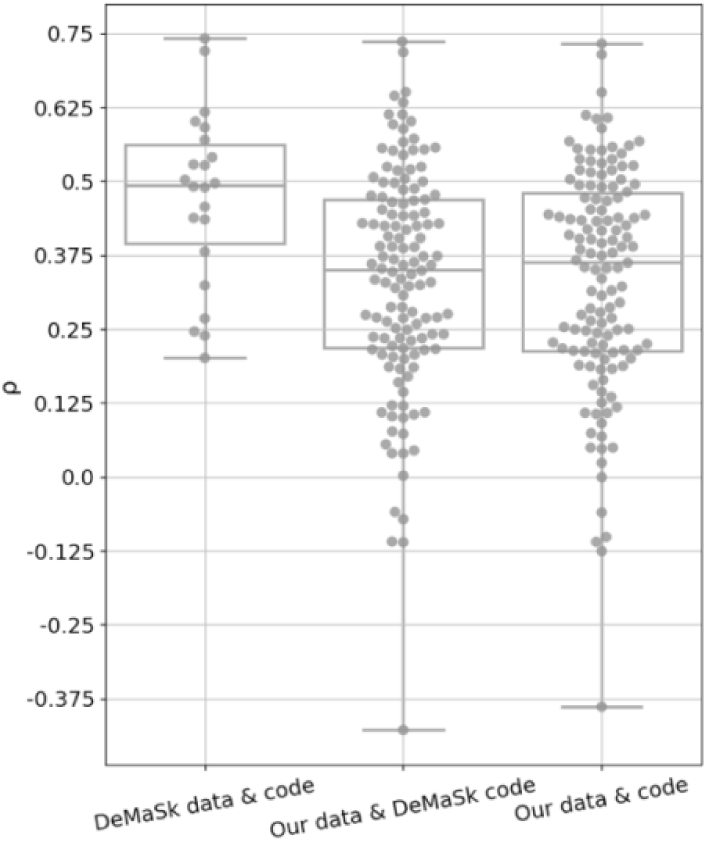
Comparison between published and re-implemented predictors. The plot shows leave-one-protein-out cross-validation performance on predictors built from the published DeMaSk code or our code. The predictors were trained and evaluated on DMS data either provided by the DeMaSk study or curated by our own. The “DeMaSk data & code” result is similar to the published result. For the “Our data & DeMaSk code” result, we used our own data and published code which shows a median performance around 0.35. This is probably because many more DMS results are included in our data. The similarity of results achieved using “Our data & code” demonstrates the correctness of our re-implementation.

**Fig S4.**
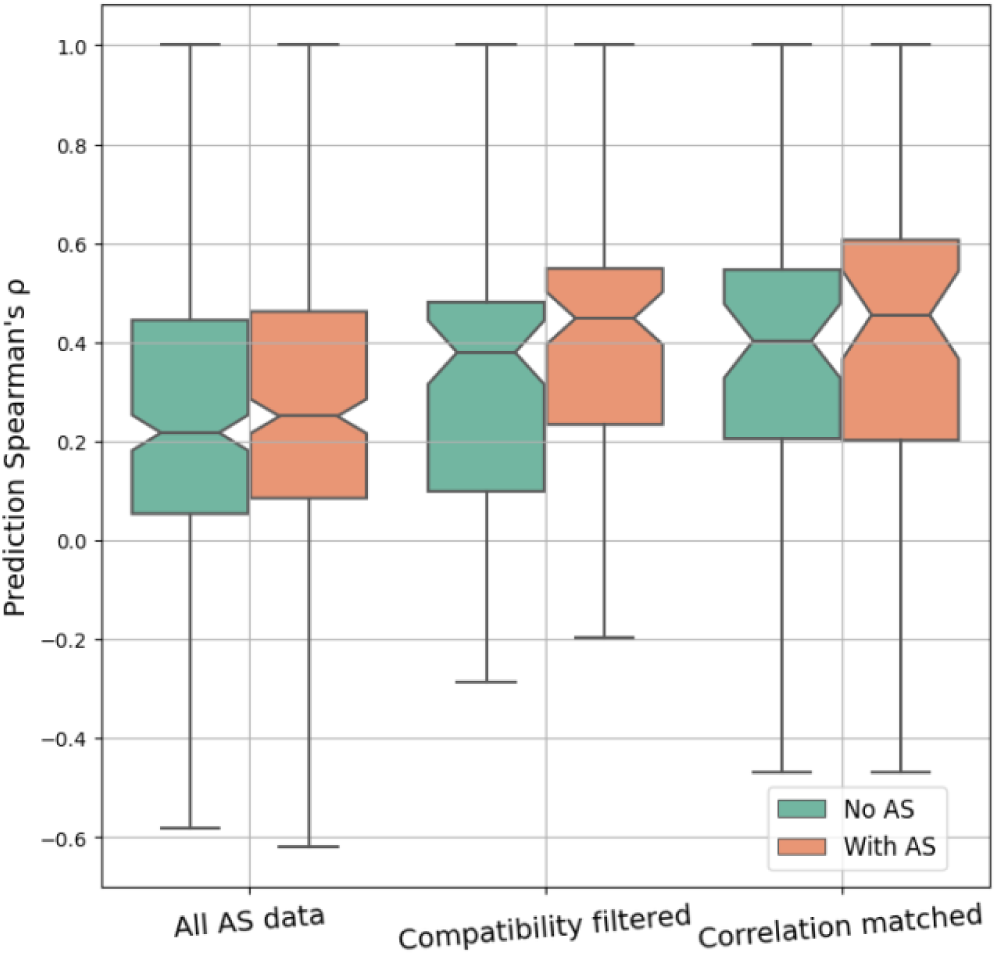
Performance comparison between predictors using AS data or not. The Spearman’s *ρ* between experiment DMS scores and predicted scores for each DMS and AS data pair are shown as box plots. Different approaches to filtering/matching the data are shown on the x-axis: “All AS data” used all available data; “Compatibility filtered” used only data of high assay compatibility; “Correlation matched” used only data with the highest regularised correlation for each DMS dataset. The figure does not include data without available (filtered/matched) AS scores. This means that the different results are not directly comparable since they are visualized on different subsets of DMS/AS data pairs (for example, “All AS data” contains all DMS/AS data pairs, but “Compatibility filtered” contains only data pairs of high assay compatibility). Control results are shown as green boxes for predicting without AS data as a feature. The underlying *ρ* for each data pair in the control results is the same, but the boxes are shifted due to data filtering/matching.

**Fig S5.**
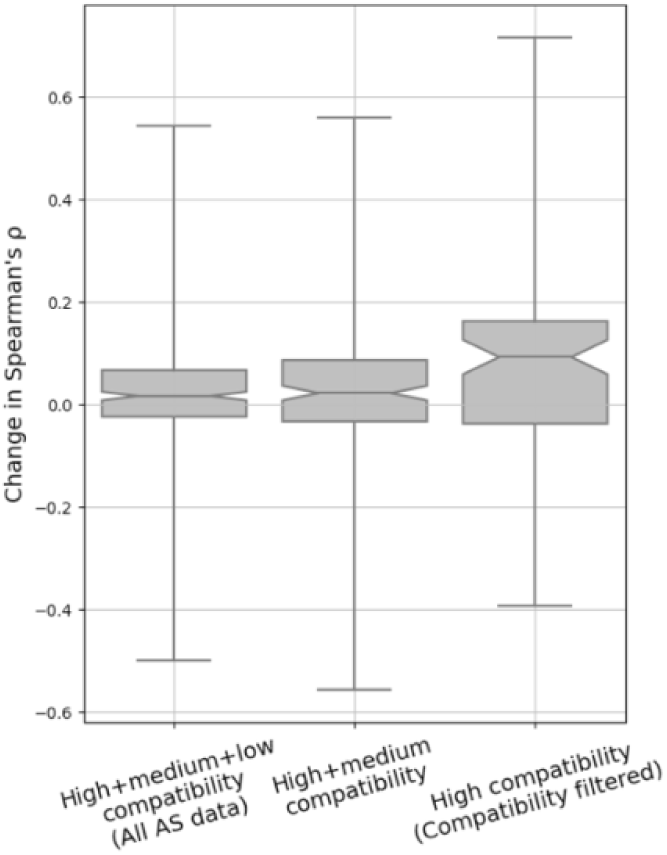
The performance of variant impact prediction for using data of different assay compatibility levels. The change of prediction Spearman’s *ρ* for each DMS and AS data pair is shown as box plots. A higher value represents higher prediction accuracy achieved for using AS data. Different data filtering methods are shown on the x-axis.

**Fig S6.**
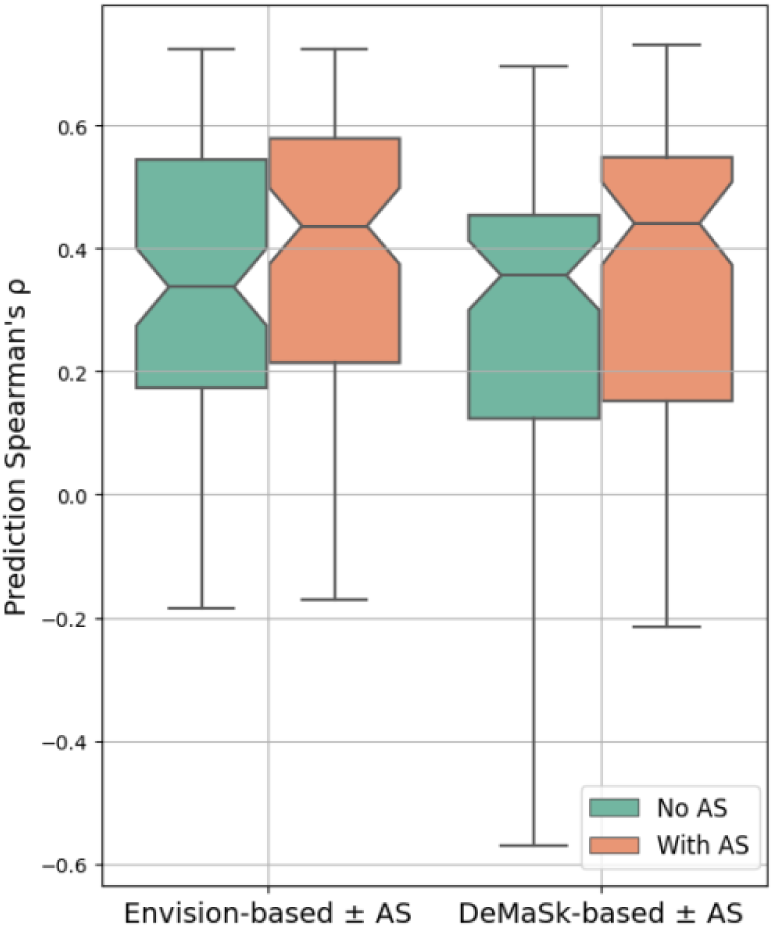
Prediction performance is improved while incorporating high compatibility AS data into the Envision model. The Spearman’s *ρ* between experiment DMS scores and predicted scores for each high compatible DMS/AS assay pair are shown as box plots. The x-axis shows the predictor used, either Envision or DeMaSk. Control results are shown as green boxes for predicting without AS data as a feature.

**Fig S7.**
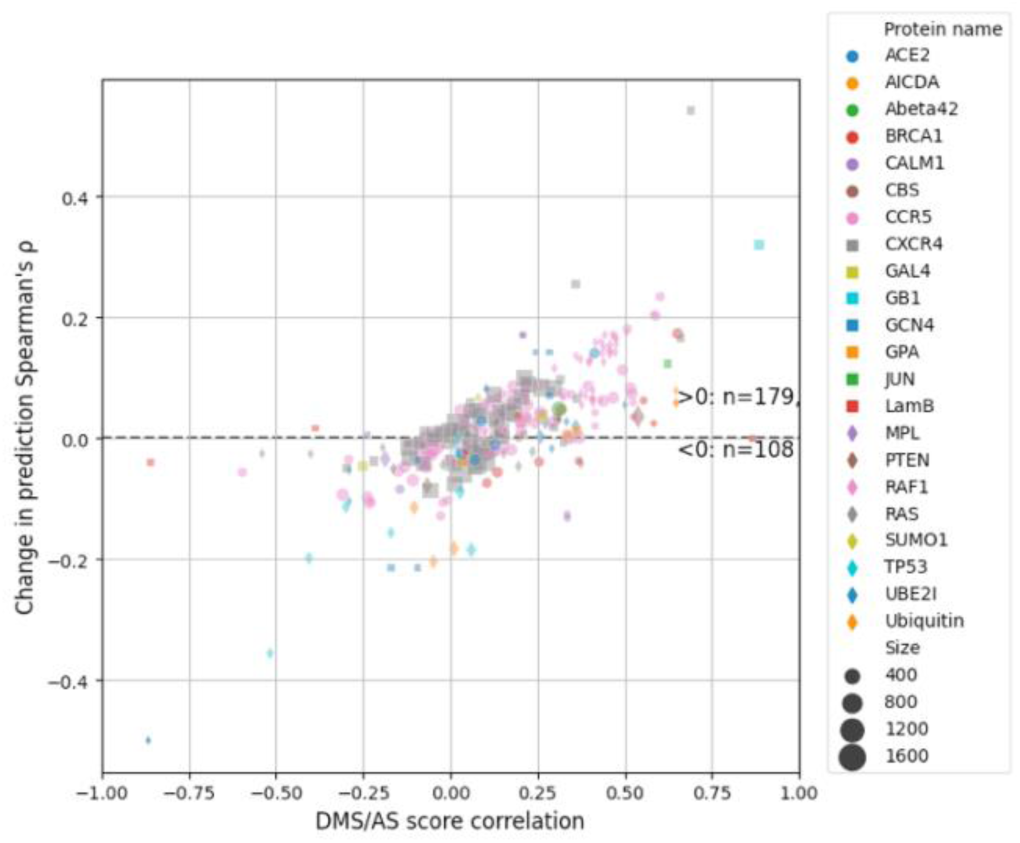
Prediction performance change for using all AS data. Each dot represents a DMS/AS data pair. The vertical axis shows the change of prediction *ρ* by using AS data (larger means higher performance achieved by using AS data). The horizontal axis shows the DMS/AS score correlation for *all* variants on the matched residues rather than just alanine substitutions. The colours and shapes of the dots correspond to the target protein, and size indicates the number of variants in each data pair.

**Fig S8.**
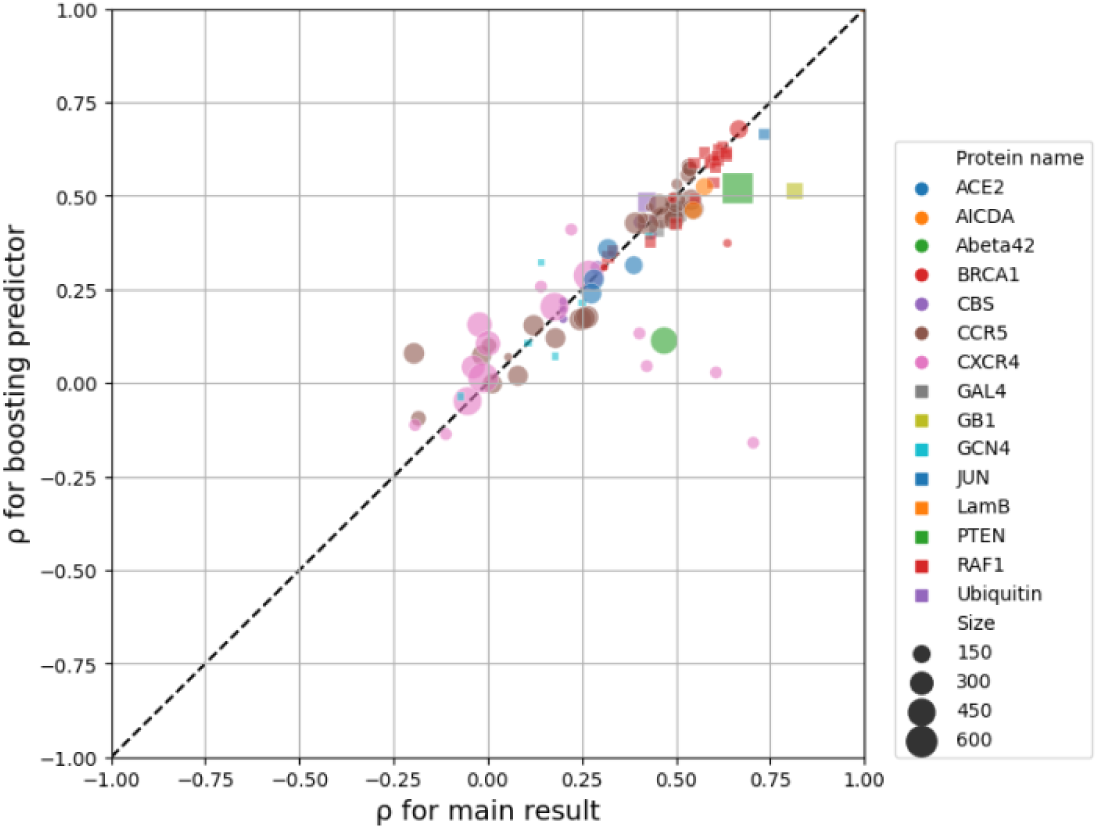
Boosting setup shows similar performance as the main result. Each dot represents a filtered DMS/AS data pair of high assay compatibility. The vertical and horizontal axes show the prediction Spearman’s *ρ* for either modelled with boosting or the one-step (main result) setup. The colours and shapes of the dots correspond to the target protein, and size indicates the number of variants in each data pair.

**Fig S9.**
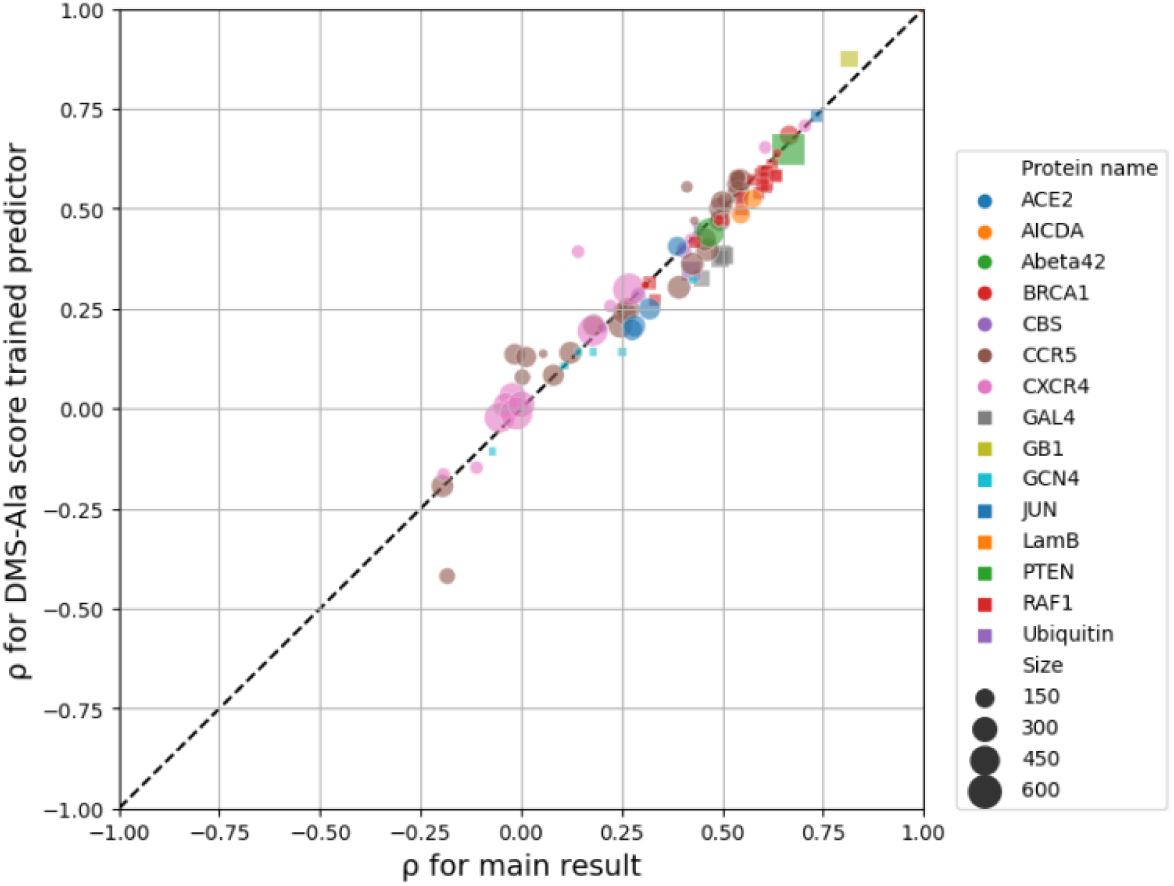
Training with DMS scores of alanine substitutions shows similar performance as the main result. The vertical and horizontal axes show the prediction Spearman’s *ρ* for predictors either trained with DMS score of alanine substitutions or AS data of high assay compatibility (main result), yet all evaluated on high compatibility AS data. The colours and shapes of the dots correspond to the target protein, and size indicates the number of variants in each data pair.

**Fig S10.**
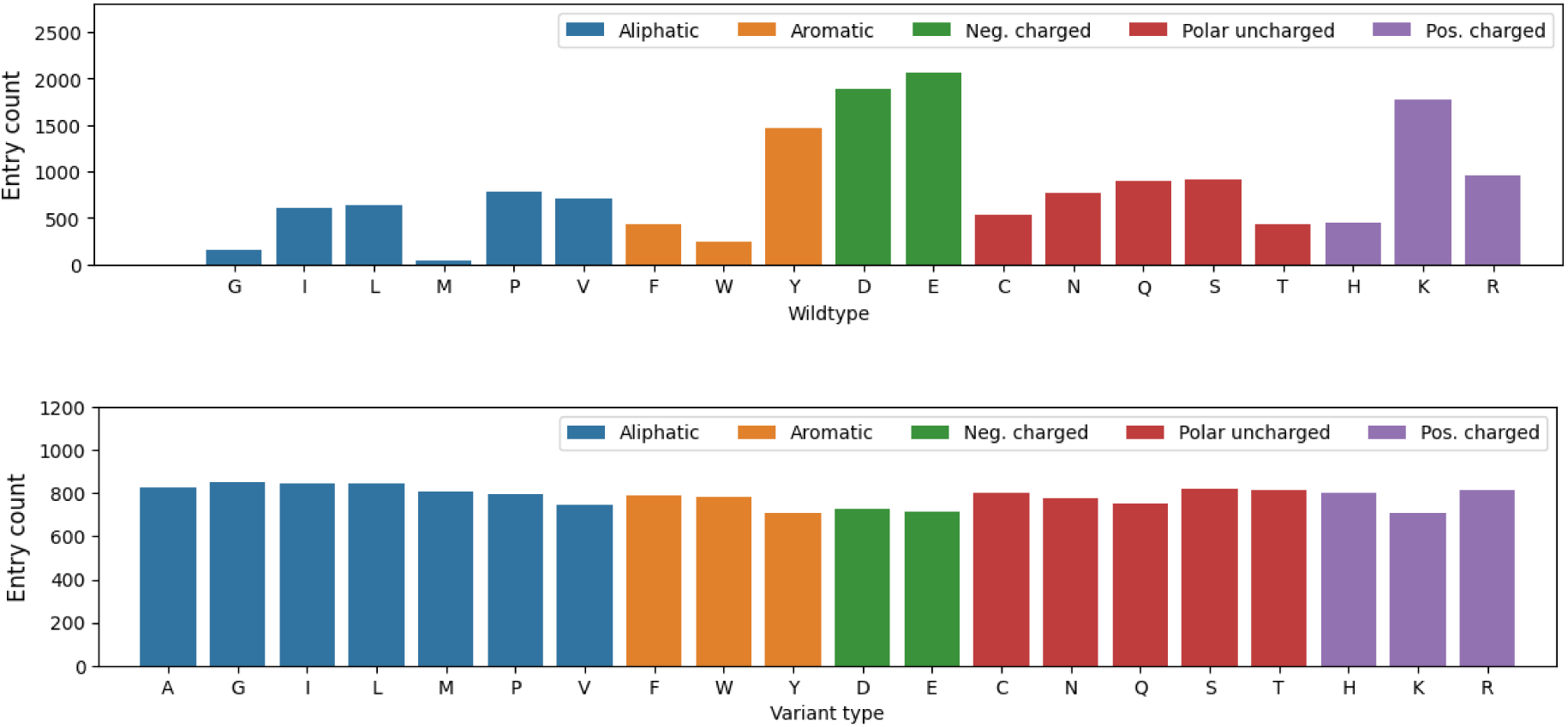
Count of variant entries for each wild-type or variant amino acid of high assay compatibility data. (Neg.: negatively, Pos.: positively)

**Table S1.**
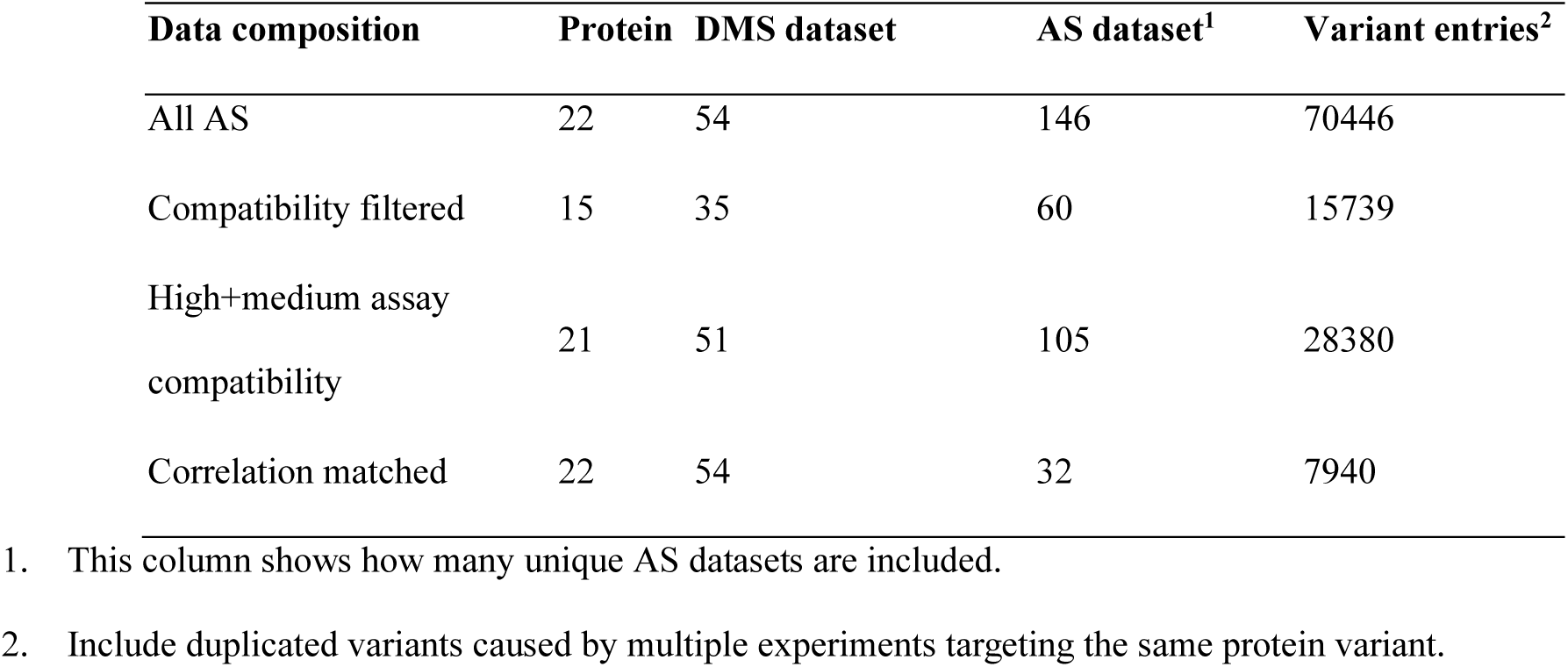
Amount of data with AS scores available

| Data composition | Protein | DMS dataset | AS dataset<sup>1</sup> | Variant entries<sup>2</sup> |
| --- | --- | --- | --- | --- |
| All AS | 22 | 54 | 146 | 70446 |
| Compatibility filtered | 15 | 35 | 60 | 15739 |
| High+medium assay compatibility | 21 | 51 | 105 | 28380 |
| Correlation matched | 22 | 54 | 32 | 7940 |

1. This column shows how many unique AS datasets are included.
2. Include duplicated variants caused by multiple experiments targeting the same protein variant.

## Supplementary information

### Applying AS data to Envision method

We re-implemented a predictor based on Envision (Gray et al., 2018) to incorporate AS data. Features used in Envision were downloaded from its online toolkit. All Envision features are used for modelling except for substitution type (wt_mut) which has low importance according to the published result and our pilot studies yet is computationally expensive in our setup. Protein data were excluded if their features were not available online. DMS and AS data pairs with high assay compatibility were used for modelling. Missing feature values were imputed by the mean values for numerical features or the most frequent values for categorical features. Categorical features are encoded with the one-hot encoder. We used sklearn.ensemble.GradientBoostingRegressor from scikit-learn package (Pedregosa et al., 2011) to build the predictor, and hyperparameters were tuned by Bayesian Optimization (González et al., 2015) with Group K-Fold (protein-30-fold) cross-validation. The training and evaluation process were similar to that previously described. For comparison, we repeated the DeMaSk-based analysis on the same subset of data.

### Boosting with AS data

To deal with the sparsity of AS data, we tested a variant impact predictor based on boosting. A first linear regression predictor was trained with all training DMS data using the three DeMaSk features without AS data, which was the same as the control predictor mentioned previously. We then calculated the prediction error by subtracting the predicted scores from DMS scores, and a second linear regression predictor was trained to predict the error. The second predictor was trained only on DMS/AS data of high assay compatibility and used both protein features and the encoded AS scores. The final prediction result was the sum of the outputs from these two predictors.

### Replacing AS data with DMS scores of alanine substitutions

We investigated another potential approach to overcome the sparsity of AS data by replacing the AS feature with the DMS scores of alanine substitutions (DMS-Ala). For all DMS datasets we collected, their AS feature values, regardless of availability, were replaced by the DMS-Ala scores on the same residue. Missing scores were imputed by the mean value of all DMS-Ala scores. A regression model was trained and evaluated as previously described, using the three DeMaSk features as well as the DMS-Ala scores. The AS data of high assay compatibility are still used for the testing process.

